# Aligning terrestrial eDNA sampling and analytical choices to effectively capture both rare species and compositional variation in grassland plant communities

**DOI:** 10.64898/2026.01.27.702028

**Authors:** Jan Plue, Mats Töpel

## Abstract

Vascular plants are a major component of terrestrial diversity, yet they are overrepresented among the world’s threatened species. To effectively manage this biodiversity crisis, data with high spatiotemporal resolution are crucial, yet often lacking for plants. Molecular assays of environmental DNA (eDNA) are capable of rapid detection of biodiversity by analyzing the collection of DNA molecules retrieved from environmental samples such as soil cores. Specifically, metabarcoding may soon generate broad-scale longitudinal plant community data, yet much methodological work on sampling strategies and analytical choices remains if soil-based metabarcoding is to become a reliable tool for monitoring both rare, target species as well as compositional changes in plant communities. Therefore, this dual-purpose study in seven Swedish semi-natural grasslands investigated if eDNA metabarcoding community data can independently 1) inform on the presence of a rare, small-statured grassland specialist (*Gentianella campestris*) and 2) simultaneously infer community compositional change. Using metabarcoding of the matK gene from the chloroplast genome, we demonstrate that eDNA metabarcoding effectively identifies a rare species in a highly taxonomically diverse habitat, uncovering *G. campestris* DNA in 31% of soil cores. Evidence suggests the eDNA signal reflects recent spatio-temporal population dynamics at fine spatial scales. Although the entire plant community was not uncovered, molecular community data proved a representative subset, effectively capturing changes in community diversity and composition at plot sizes commonly used for plant surveys. Choices surrounding relative read abundance filtering had significant bearing on eDNA’s discriminating power: filtering may conservatively remove true observations of a rare species, while filtering highly localized plot noise led to more robust patterns emerging in species richness and plant composition turn-over. Given careful alignment of study goals and sampling strategies, soil-based eDNA metabarcoding may already provide a stand-alone tool for generating reliable, scalable and observer-independent longitudinal data for monitoring changes in terrestrial plant communities.

## Introduction

Vascular plants are a major component of terrestrial biomass and diversity (Bar-On et al. 2018; Kier et al. 2005), being crucial for biodiversity at higher and lower trophic levels (Wan et al. 2020), ecosystem processes (e.g. carbon sequestration, Chen et al. 2020), ecosystem stability (Haddad et al. 2011) and ecosystem service provisioning (Letourneau et al. 2011). At the same time, plants are clearly overrepresented among species at risk of extinction (ca. 40% of assessed species across all taxa, Nic Lughadha et al. 2020). A range of global change drivers such as habitat loss, climate change and eutrophication are rapidly transforming and/or eroding plant community diversity at local, regional and global scales (Vellend et al. 2013, Nic Lughadha et al. 2020, Kimmel et al. 2022, Montràs-Janer et al. 2024), underpinning repeated, urgent calls to hasten and improve upon habitat restoration and conservation efforts (Banks-Leite et al. 2020, Watson et al. 2016). However, to identify and effectively manage biodiversity during this conservation crisis, access to monitoring data with high temporal and spatial resolution that can accurately inform on the presence of rare, threatened species and compositional community change is crucial (Rodrigues et al. 2006). Still, such effective dual-purpose monitoring programs are mostly lacking in vascular plants (Bachman et al. 2019, Jeliazkov et al. 2022).

Environmental DNA (eDNA) refers to genetic material recovered directly from any type of environmental or biological sample, which can subsequently be analyzed using molecular approaches to assess biodiversity through the identification of species-specific DNA sequences (Thomsen & Willerslev 2015). Molecular analyses of eDNA may thus offer considerable potential for generating broad-scale longitudinal data for terrestrial plant biodiversity monitoring (Yoccoz et al. 2012), particularly as terrestrial plant eDNA applications, both species-specific assays as well as metabarcoding, have shown great promise in a range of habitats (tundra, Edwards et al. 2018; desert, Carrasco-Puga et al. 2021; grassland, Hartvig et al. 2021; boreal forest, Ariza et al. 2023; mediterranean and tropical communities, Duley et al. 2023). However, species-specific molecular assays - while potentially offering higher sensitivity for detecting individual, rare species - cannot inform on plant community composition. By contrast, soil metabarcoding offers the possibility of simultaneously detecting rare target species and characterizing broader plant community composition within a single workflow. Its value for conservation and restoration applications therefore lies precisely in this potential dual use. Still, whether broad metabarcoding markers can reliably recover ecologically meaningful signals for both rare species occurrence and community composition remains poorly resolved. While being a potential key tool of future plant community research and monitoring, much field-, lab- and bioinformatic methodological development thus remains if terrestrial eDNA metabarcoding is to become a reliable stand-alone tool to perhaps one day replace traditional plant community surveys otherwise hampered by high human-resource and monetary costs, temporal sampling constraints (Yoccoz et al. 2012) and observer bias (Verheyen et al. 2018).

Any observational study or monitoring scheme is only as good as its sampling design and the degree to which samples are representative of the target ecosystem, both of which are ultimately guided by the study or monitoring objectives. In other words, how well an individual sample reflects the local extant presence of individual species and/or community composition, will determine the chances of detecting subtle spatio-temporal changes in species distributions or community composition across multiple locations. Sampling invisible, below-ground phenomena, *in casu* DNA distributed in the soil, therefore presents a formidable challenge when the aim is to acquire an accurate and representative local sample (cf. Baker et al. 2009; Muukkonen et al. 2009; Plue & Hermy 2012). This issue must be properly addressed during soil eDNA sampling for metabarcoding applications if firm ecological conclusions are to be drawn from resulting eDNA data on either individual species occurrences or plant community composition (Zinger et al. 2019). The challenges for representative sampling are indeed numerous. First, established plants are often spatially clustered in response to environmental heterogeneity (Ulrich et al. 2021), which may trigger similarly clustered belowground patterns in roots, seeds, pollen and associated intra- and extracellular DNA (Pärtel & Wilson 2002; Plue & Hermy 2012; Pardoe 1996; Levy-Booth et al. 2007). The resulting high spatial heterogeneity in soil DNA therefore presents a significant obstacle when extracting a single plot-based sample that is expected to represent the local, extant plant community. Second, plant soil eDNA may contain and accumulate DNA originating from foreign sources that are spatially or temporally disconnected from the current local plant community (Levy-Booth et al. 2007). Non-local DNA may originate from far-reaching roots (Hiiesalu et al. 2012), pollen dispersal (de Vries et al. 2003) or extracellular DNA transported within the soil matrix (Poté et al. 2007). In contrast, so-called legacy DNA refers to DNA persisting from past historical inputs from plants no longer present locally, such as temporally dispersed seeds (Plue & Cousins 2018), persistent intercellular DNA from leaf and root litter or extracellular DNA adhered to soil complexes (Levy-Booth et al. 2007, Yoccoz et al. 2012, Carini et al. 2016, Ariza et al. 2023). Each of these processes may, on its own or in combination, reduce the probability of acquiring a reliable local, plot-based signal, and may thereby distort true plant community patterns ((cf. Carini et al. 2016; but see Edwards et al. 2018; Ariza et al. 2022 for high eDNA–community taxonomic overlap)). Still, such legacy or non-local signals need not be purely problematic. Indeed, whether such, often low-abundant, signals should be interpreted as noise, bias or ecologically meaningful information (e.g. wider species pool or restoration potential) depends strongly on the objectives of the study and on how low-abundance or non-local detections are treated in subsequent analyses.

An added challenge to a single, dual-purpose eDNA metabarcoding workflow is indeed that eDNA metabarcoding data are not only shaped by the above mentioned ecological processes, but are further susceptible to methodological, technological, analytical and bioinformatic biases. Low sequence read abundances are therefore often seen as problematic and difficult to interpret as they may reflect 1) true detection of a rare species, 2) poor amplification of an abundant species (Elbrecht & Leese 20153), 3) low quantities of non-local DNA, 4) variation in sequencing depth (Deagle et al. 2019) or 5) stochasticity during (sub)sampling, DNA extractions, PCRs and DNA-sequencing (Luo et al. 2023). This perceived risk of field- or lab-contamination producing false positives thus often translates into low-abundance reads being conservatively removed. This creates an important trade-off for conservation applications: filtering may improve the robustness of community-level patterns by reducing noise, but may also precisely remove the low-read signals that are most valuable to trace rare, threatened species (Hartvig et al. 2021), detect the onset of species invasions (Sepp et al. 2021), evaluate a site’s restoration potential or assess the plant community beyond the spatial and temporal limits of the local plot sample (cf. Carrasco-Puga et al. 2021). This trade-off may be particularly pronounced in species-rich plant communities (with most eDNA plant community studies have focused on species-poor communities (Edwards et al. 2018; Ariza et al. 2023)), where the majority of plant species are naturally low abundant (Ulrich et al. 2010), and therefore the amount of eDNA they shed and thus their detection likelihood also is low (Yoccoz et al. 2012; Edwards et al. 2018). Consequently, the usefulness of soil metabarcoding as a surveying tool depends not only on whether plant DNA can be detected, but on whether sampling design and post-hoc data curation can be aligned within the specific inferential goal: detecting rare, target species, characterizing and detecting changes in plant community composition, or doing both simultaneously.

Taken together, translating soil-derived DNA sequence reads into plant species occurrence and plant community data remains challenging, particularly when the aim is to retain a representative and reliable local plot-level signal. Yet, this challenge is precisely where soil metabarcoding may be most valuable for monitoring for conservation and restoration purposes: the same community-level assay could, in theory, detect both rare, target species of conservation interest while providing valuable information on broader spatio-temporal community patterns. To test this potential, we designed a dual-purpose study in seven species-rich semi-natural grassland communities in Sweden, each containing current or historical populations of the rare grassland specialist species *Gentianella campestris* L. We selected *G. campestris* as a deliberately challenging test-case because the species is small-statured, rare and biennial, and is therefore expected to contribute only limited amounts of DNA to the soil. If such as a species can be detected using broad metabarcoding while the same samples also recovers meaningful community-level variation in highly diverse semi-natural grassland communities (Wilson et al. 2012), this would provide a strong case that soil eDNA metabarcoding, as opposed to single-species assays, may be uniquely suited for monitoring plant communities for conservation and restoration purposes. Specifically, we therefore questioned: 1) under which sampling conditions can soil eDNA metabarcoding successfully detect *G. campestris* DNA, 2) at what plot size is the plant community most accurately represented by a single eDNA soil core, 3) at what plot size do eDNA-derived plant community data capture plant compositional differences between plots and 4) how does post-hoc removal of low abundance reads affect rare-species detection and community-level inference?

## Materials and Methods

### 2.1 Study area

We performed the study in seven managed semi-natural grasslands scattered across northern Stockholm county in central Sweden. Nordic semi-natural grasslands are known for their exceptionally high species diversity at small scales due to a long history of continuous grassland management (0.1 – 1 m^2^; Wilson et al. 2012). Because of severe global declines (e.g. Auffret et al. 2018), semi-natural grasslands are today of global conservation concern (Watson et al. 2016) triggering ongoing efforts to monitor their biodiversity over time (National Inventory of Landscapes in Sweden, Ståhl et al. 2010; National Meadow and Pasture Inventory, http://www.sjv.se/tuva), rendering them particularly suitable for our study’s main goals. All grasslands (Table S1), ranging in sizes between 4011 and 128 423 m^2^ (mean of 45 554 ± 15 516 m^2^) were managed by either mowing (two sites) or livestock grazing by cattle (three sites) or sheep (two sites). A final selection criteria for the study sites was the recorded and geo-located presence of the rare, red-listed grassland specialist species *Gentianella campestris*, with selected grasslands representing a gradient in the time since *G. campestris* was last observed and a gradient in population size.

### 2.2 Data collection

#### eDNA samples

During October 2021, we collected individual soil cores with a sampling scheme (cf. Fig. 1) which would help us (1) identify the sensitivity of eDNA at detecting *G. campestris*, and how sampling and population factors affect that sensitivity and (2) enable us to investigate if and at what spatial scales (i.e. those generally applied during traditional plant community inventories) eDNA core samples would effectively capture variation in plant community and diversity. Based on available GPS coordinates extracted from the Swedish Species Observation System (Artportalen; www.artportalen.se), we located where *G. campestris* was last observed within each grassland. We then visited these coordinates and collected a total of nine soil cores from each grassland, i.e. three samples randomly spread out in each of three circular bands around these coordinates (0 – 5 m, 5 – 40 m, 40 – 120 m). The coordinates of each sampling point were documented using a high performance GPS (Emlid Reach RS2 RTK) to an accuracy of 10 cm. The soil auger (3.5 cm diameter) was thoroughly cleaned between each soil core sampling event. First, we used a dish brush with de-ionized water to clean the auger of any visible soil and plant material. The auger was then rinsed profusely twice, first with de-ionized water followed by a 5% chlorine solution, the latter to destroy any remaining DNA. A mineral soil core (i.e. removing the litter layer and plant parts prior to sampling, using single-use medical gloves) was then collected to a depth of 5 cm, and placed directly into a plastic zip-lock bag. Soil core samples were immediately put in a cool box and subsequently in a −21°C freezer at the end of each day of fieldwork, ensuring minimal impact on eDNA sample quality (Frøslev et al. 2023).

**Fig. 1.**
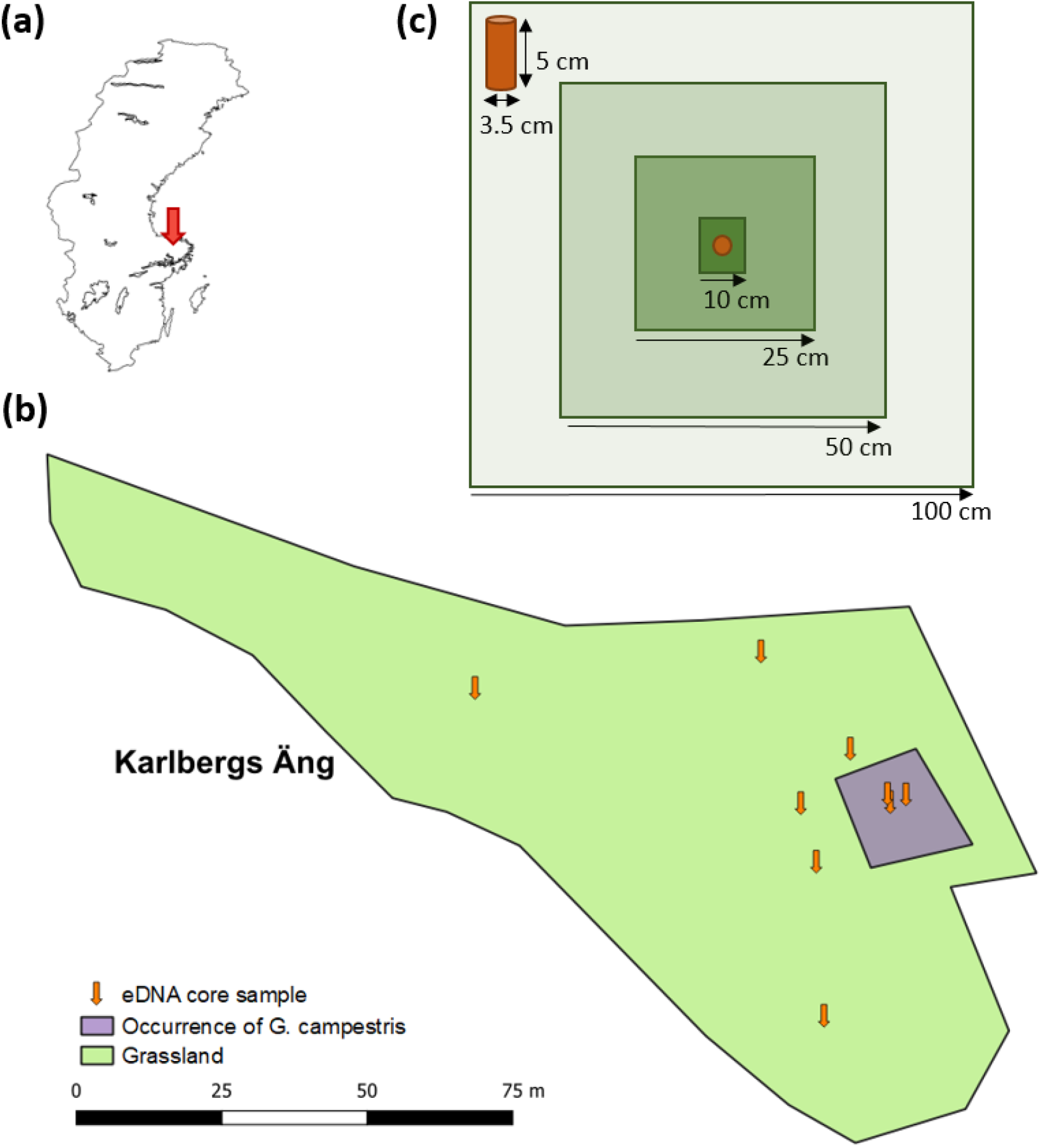
eDNA sampling design applied across 7 managed semi-natural grasslands in Stockholm county, Sweden (a). Nine eDNA soil core samples (gold arrows) were collected per grassland (b), at increasing distances away from the centre of occurrence of *Gentianella campestris* (purple polygon) in the grassland (green polygon). Around each core sample, plant species presence was recorded in four plots of increasing sizes (10, 25, 50 and 100 cm) centred around the soil core (c). The sampling design aimed to investigate terrestrial eDNA detection likelihood of a rare grassland plant specialist and eDNA’s potential to representatively capture variation in plant community composition.

#### Sampling and population information

While collecting soil cores in 2021, we equally recorded the distance from each soil core to the nearest *G. campestris* individual. At the same time, we counted all *G. campstris* individuals in a 2 m radius from the soil core to estimate 2021 population density. Upon returning for traditional plant community inventories in 2022, we again counted all *G. campestris* individuals within a 2 m radius of each core sample location to estimate the 2022 population density. Finally, we extracted population size data from the Swedish Species Observation System, where designated flora guards (SE Floraväktarna; <u>Website</u>) regularly report population sizes of red-listed species from locations to which they were assigned. As such, we have exact data on *G. campestris* population size during the last eight years, as well as information on the year when the species was last visually observed. As the species is biennial with significant inter-annual demographical variation (Lennartsson & Oostermeijer 2001), we used the eight year population size average as a population size proxy.

#### Plant community inventories

In June 2022, we used the same high accuracy GPS (Emlid Reach RS2 RTK) to return to each of the 63 sampling locations. For 62 samples, we managed to re-locate the exact hole which was left behind after autumn core sampling. We inventoried plant communities centered around the core sample location along a nested plot design. Plot size increased from 10 × 10, to 25 × 25, 50 × 50 and 100 × 100 cm^2^ centered around the soil core location. Per plot size, we recorded the presence and cover [%] of all vascular plant species, returning a total of 252 plant community inventories.

### 2.3 DNA extraction and PCR

Soil samples were transferred to 50 ml Falcon tubes and then freeze dried for approximately 24 hours. Glass beads were then added and the samples were homogenized using a FastPrep-24 5G (MP Biomedicals, LLC). DNA was extracted using the DNAeasy power soil pro kit (Qiagen) according to the manufacturer’s instructions. DNA-concentration was then quantified using Qubit™ 1X dsDNA High Sensitivity (HS) assay kit in the Qubit Flex fluorometer (Thermofisher). DNA concentration of each sample was diluted to 10 ng/µl after which the multiplex PCR reactions were performed in 25µl volumes containing 2µl template DNA, 12.5 µl 2X Phusion™ Plus Green PCR Master Mix, 1.25 µl of 10 µM forward and reverse primer solution mixes (see Table 1) and 8 µl DNase free water. The PCR-program started with 2 min. denaturation at 98°C followed by 5 cycles of 10 min. at 98 °C, 35 sec. at 53 °C and 30 sec. at 72 °C, then 35 cycles of 10 sec, at 98 °C, 30 sec. at 63.6 °C and 30 sec. at 72 °C and finalised with a 5 min. extension period at 72 °C. Each sample was amplified in triplicate and then pooled before being purified using the AMPure magnetic beads assay (Beckman Coulter, US). The PCR also included three control samples, one negative with DNase free water instead of DNA-template, and two positive with 2µl template DNA from *Helianthemum nummularium and Elytrigia* repens, respectively.

**Table 1.**
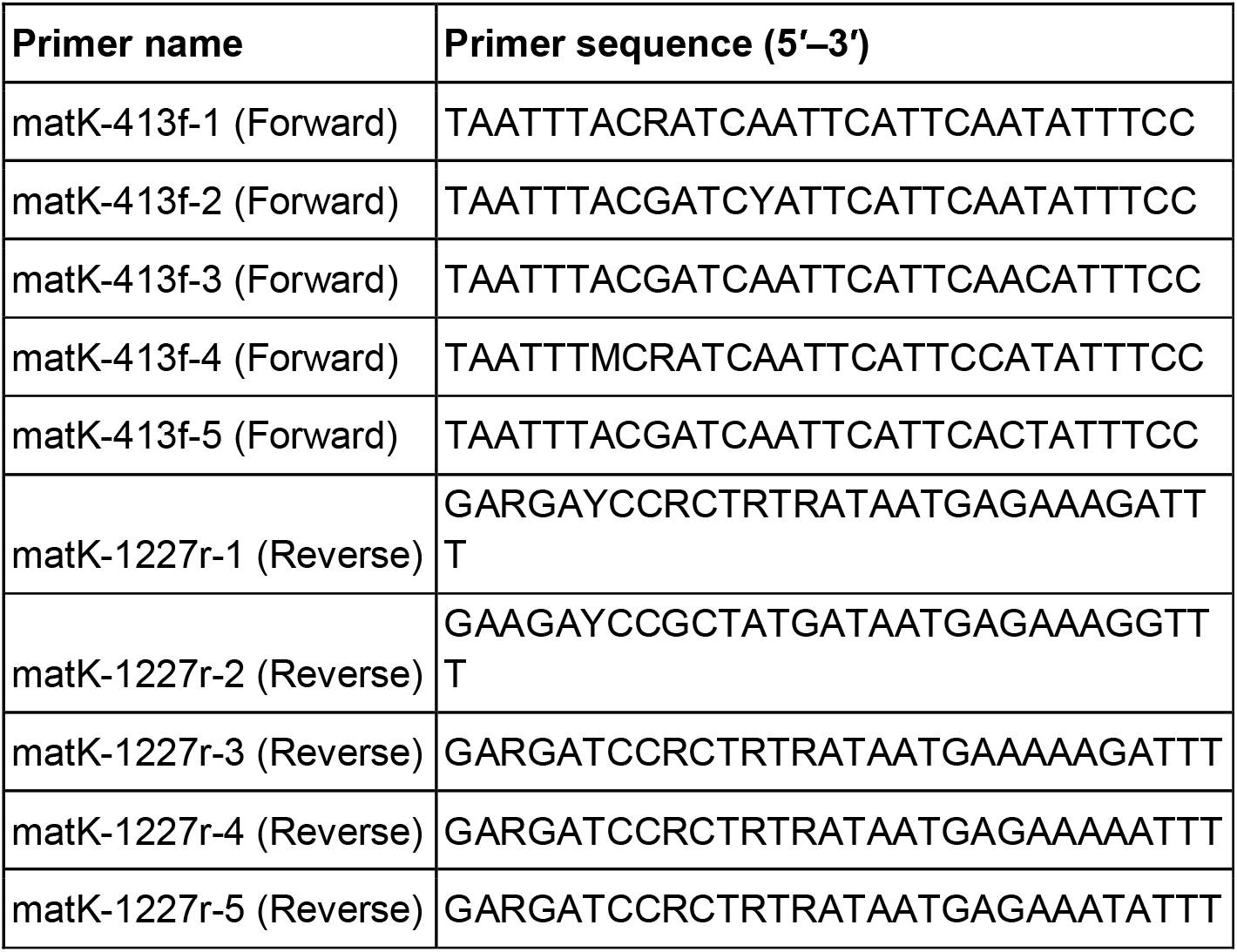
The partially degraded primer sequenced from Heckenhauer et al. (2016) used to amplify the matK region of the chloroplast.

A second PCR was performed in a final volume of 25 μl containing 2 μl of purified PCR-product, 5 μl of 5X Q5 reaction buffer, 0.4 mM M13 forward and M13 Reverse index primers, 0.2 mM dNTPs, and 0.025 U/μl Q5 High-Fidelity DNA Polymerase (New England Biolabs), following the PCR-protocol provided by the sequencing facility SciLifeLab, Uppsala. Amplicons were then purified with SMRTbell shape cleanup beads to remove too short fragments.

### 2.4 Bioinformatic analysis and quality control

The DNA-library was sequenced on one SMRT® Cell 8M v3 of the PacBio Sequel II platform at the Uppsala Genome Center of the National Genomic Infrastructure, SciLifeLab. The resulting sequences were processed using the nf-core/ampliseq pipeline v2.7.0 (Ewels et al. 2020), that included quality analysis using fastQC (Andrews 2010), trimming and adaptor removal using cutadapt (Martin 2011), sequencing error correction and taxonomic classification of the amplified sequence variants (ASVs) using *dada2* (see Table S2 in Supplementary Materials for parameter settings; Callahan et al. 2016). For the taxonomic classification, a custom reference sequence database was used containing 4946 sequences downloaded from the National Center for Biotechnology Infrastructure (https://www.ncbi.nlm.nih.gov/) from 2516 species known to occur in Sweden.

The number of raw reads for each sample ranged between 30450 - 105827 sequences (median 54527) with the exception of four samples with low yields (1100, 121, 79 and 67 reads respectively). These low yield samples were not excluded from the analysis as they did not behave as outliers by any other measure amongst the derived variables, and their omission had no effect on the results. Between 41-83% of the reads remained after quality filtering and adaptor trimming and were subject to taxonomic classification by dada2 (Callahan et al. 2016). In total, 731 ASVs with a median length of 828 bp (min. 791 bp, max. 928 bp) were detected in the 62 soil core samples. Some 622 were taxonomically classified to species level and one only to the genus *Convallaria*. The remaining ASVs were identified as *Streptophyta* (78) or as *Eukaryota* (29). In addition, 69 ASVs had an exact match (via *species_exact* by dada2) to a sequence in the reference database.

Then, we manually curated the raw ASV data for quality control and data cleaning purposes. ASV nucleotide sequences were subject to similarity searches using BLASTn (via BLAST+ v2.14.0; Camacho et al. 2009), querying sequences against the NCBI nucleotide database using default parameters. Following this, a series of changes were made to the automatic bioinformatic classification. First, all ASVs previously only identified as *Streptophyta* could be assigned to the species (71) or genus level (7). Most ASVs were of species already identified via the bioinformatic pipeline, but 13 ASVs were assigned to nine species previously unidentified by the bioinformatic pipeline (*Galium mollugo, Geum rivale, Myosotis stricta, Ranunculus acris, R. repens, Saxifraga granulata, Sesleria caerulea, Solidago virgaurea* and *Stellaria alsine*). Second, ASVs only classified as *Eukaryota* proved to be from bacteria and were omitted. Third, species assignments to reference sequences representative of multiple congeneric species were assigned to a single species based on either BLASTn assignment or expert judgement based on Swedish species distributions or clear differences in habitat preferences. For example, an ASV assigned to reference sequence KF737610.1 representing both *Sagina procumbens* and *S. saginoides* was identified as *S. procumbens* since *S. saginoides* most southern Swedish population was ca. 400 km to the east of our study grasslands. Fourth, BLASTn identified that two species (*Festuca rubra* and *Galium boreale*) could be further subdivided into *Festuca rubra, F. ovina, Galium boreale* and *G. verum*. Our study thus clearly shows that long read sequencing of soil eDNA targeting the current plant community is both feasible and able to result in a high proportion of species-level classifications (88% from automatic classification, and 98% when adding manual classification), in part likely due to using only a reference database limited to local (Swedish) plants (cf. Barnes et al. 2022).

Finally, the three control samples were used to clean the data matrix further. The two positive control samples generated in total 48096 and 41102 reads, respectively. Of these, the majority originated from the reference species *Helianthemum nummularium* and *Elytrigia* repens, as expected. Low read counts of the high concentration reference template DNA of either species (totals read counts of 38 and 52, respectively) were recorded across 5 and 7 field samples, respectively. Then, two ASVs were additionally uncovered in the positive controls, classified as *Plantago lanceolata* and *Galium boreale* and represented by 29 and 13 reads, respectively. In the negative control sample, 68 reads were generated, i.e. 5 from *H. nummularium*, 16 from *E. repens*, 16 from *P. lanceolata*, 9 from *G. boreale* and another 22 reads from two ASVs unique to the negative control sample, identified as *Agrostis capillaris* (20) and *Vaccinium vitis-idaea* (2). The positive and negative controls thus indicate that minor contamination between field samples may likely have occurred during the laboratory analysis. In response, the reference species as well as the unique ASVs of *Agrostis capillaris* and *Vaccinium vitis-idaea* were removed from the dataset. For *P. lanceolata* and *G. boreale*, the low read counts did not justify removing the species entirely, representing 0.007 and 0.008% of their total read count, respectively (given them being the third and fourth most common species found in the field samples; cf. Dataset S3). Therefore, these species were retained in the final dataset. Instead, we relied on the independently established 0.05% RRA threshold (cf. Deagle et al. 2018) to reduce potential low-level contamination (≤ 0.05%), including traces from species both detected and not detected in the controls, while retaining low-abundance species detections in these highly diverse grassland communities (> 0.05% RRA) as true occurrences. The positive and negative control samples were thus not used to determine the RRA threshold. Both the ASVs and the final, manually curated 67 species × 63 plot data matrix are available on Dryad (Anonymous 2026).

### 2.5 Data analysis

#### Data preparation

The primary objective of the study was to evaluate if data on the presence-absence of a rare species (*G. campestris*) as well as data on the plant community unveiled via eDNA metabarcoding can be used as a stand alone information source to accurately inform on plot-based species detection and plant community composition in the established vegetation, inferring species distributional and community compositional changes between plots. To investigate if mitigation of methodological, technological and analytical bias in an ASV data matrix is dependent upon study objectives, we used both a full and a reduced species × sample ASV data matrix (See Datasets S2 and S3; will be deposited at Dryad). The reduced ASV data matrix was calculated from the full data matrix. We used sample-based relative read abundances (RRA) to remove all ASVs detected below a threshold of 0.05% of the total number of ASVs for that sample (Deagle et al. 2019), hypothesizing that this would 1) decrease the likelihood of detecting rare species and 2) increase the likelihood that eDNA plant community data would be a more accurate representation of the established plant community and compositional trends among plots. Finally, both the full and reduced ASV data matrices had ASV read abundances transformed to presence-absence.

#### Detection likelihood

To investigate if the chance of detecting *G. campestris* DNA was dependent upon sampling and inter-annual variation in population characteristics, the presence/absence of *G. campestris* in the 63 eDNA soil samples was analyzed using mixed effect logistic regressions (*glmer* from *nlme* package, Binomial distribution with *logit* link function). We constructed a full model focusing on testing the combined impact of sampling distance and population characteristics:

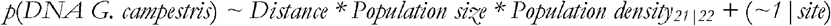

This model contained the main variables of interest as fixed effects and potential interactions, while grassland site was incorporated as a random intercept to accommodate for confirmed inter-site variation (likelihood ratio Chi^2^ = 5.54; p = 0.02; Zuur et al. 2009). However, given high collinearity between distance and population density_21|22_ (Spearman rho_21|22_ = −0.65 | −0.61, p < 0.001), we could not reliably combine these variables in a single model. Instead, we added each fixed effect separately to a null model containing only the random intercept term, using likelihood ratio tests (function *anova*) to identify the direct importance of distance, population size and population density (for both 2021 and 2022) separately. In a next step, we tested the additive effect of population size on detection likelihood, by adding this predictor variable to the random effect (*site*) models containing either distance or population density (only 2021) as a fixed effect, as these predictor variables did not co-vary strongly with population size (Spearman rho = −0.24, p = 0.06; Spearman rho = 0.31, p = 0.01, respectively). Two-way interactions in these models were tested but proved non-significant. All predictor variables were scaled so that comparable standardized effect sizes (SES) would be returned (Grueber et al. 2011), which allows for comparison of effect sizes of each predictor within and across models. Finally, to investigate the effectiveness of eDNA to trace a rare species and the impact of data curation on sampling effort guidelines for its detection, the above analysis was run on two presence/absence variables of *G. campestris*: 1) using all recorded presences, irrespective of the number of detected ASVs and 2) using only those records with an RRA > 0.05%.

#### Community representation

To test if, and at what plot size, species richness captured via a single eDNA soil sample differs from the species richness from the vegetation, we calculated species richness per sampling effort (eDNA core sample location = 0, 10 cm distance, 25 cm, 50 cm and 100 cm). Species richness was then analyzed using generalized linear mixed effect models following:

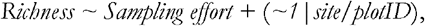

including *sampling effort* (5 factor levels) as fixed effects and *plotID* nested in *grassland site* as a random intercept, with a Poisson error distribution, respectively (using the *glmer* function in the *nlme* R-package). *plotID* was included to account for the longitudinal, i.e. nested, plant community inventories centered around each soil core. Homogeneous subsets were identified by resetting the base factor level for *sampling effort* used for comparison and re-running the model.

Furthermore, to investigate if, and at what plot size, a single eDNA soil sample may most accurately represent extant plant community composition, we calculated the Raup-Crick similarity metric (*vegdist* function in the *vegan* R-package; Oksanen et al. 2023) to assess compositional similarity between the plant community identified in an individual soil sample using eDNA vs. the plant communities inventoried at various plot sizes. For comparison to other studies, we also calculated the Bray-Curtis similarity metric which is prone to severe bias when species richness differs between communities under comparison (Koleff et al. 2003; as observed in this study). Bray-Curtis similarity generally overestimates similarity, a problem avoided via random draws from the entire species pool, rendering Raup-Crick similarity less sensitive to species richness bias and therefore ought to be the preferred metric (Vellend et al. 2007). We used a general linear mixed effect model to compare whether Bray-Curtis similarities were consistently larger than Raup-Crick similarities with *metric* (Bray-Curtis vs. Raup-Crick) as fixed effect and *plotID* (to account for longitudinal sampling) nested in *grassland site* as a random intercept, with a Gaussian error distribution (*glmer* function in the *nlme* R-package).

In order to understand which species may drive observed changing patterns in compositional similarity with plot size, we then calculated three species richness variables per plot size: 1) the number of species unique to the traditional survey data, 2) the number of species unique to the eDNA data and 3) the number of species shared by the eDNA- and traditional survey data. These variables were analyzed following the same generalized linear mixed effect model structure as for species richness, with main differences being 1) *sample effort* comprising only 4 factor levels representing different plot sizes and 2) the use of Gaussian error distribution for the similarity metrics (square root-transformed for normality). Finally, to assess the impact of data curation on potential patterns of representativeness, the above analyses were rerun using variables calculated based on the species × sample presence/absence matrix in which all RRAs < 0.05% had been removed.

#### Capturing and predicting plant community variation

We finally needed to establish whether eDNA-based plant community data can be reliably used as an accurate, stand-alone indicator of the true spatio-environmental variation in species richness and community composition. Therefore, we first ran general linear mixed models per plot size (including *grassland site* as a random intercept) to test if and at what plot size eDNA species richness would predict species richness observed in the vegetation. Models were run separately with eDNA species richness based on either the full or 0.05% RRA filtered species × sample matrix to assess how data curation by removing ASVs of low abundance affects discovery in species richness prediction. Secondly, to assert if eDNA plant community data can detect spatio-environmental variation in observed plant community composition we applied non-metric multidimensional scaling to each of five full presence/absence species × sample matrices (one per sampling effort: eDNA and four plot sizes). For reasons outlined above, Raup-Crick similarities were chosen as the distance metric underpinning the NMDS, whereas we constrained the optimal solution to returning only two axes, rotated to present the main axes of compositional variation in each dataset. For each NMDS, we then extracted the axis scores for both axes for each of the 63 samples. To check whether compositional variation was similar in paired vegetation and eDNA communities, general linear models were then built per plot size where eDNA and vegetation composition (NMDS axes) were the response and predictor variable, respectively. The entire compositional and post-hoc analysis outlined before were subsequently rerun based on the 0.05% RRA filtered species × sample matrix to assess how data curation affects compositional pattern discovery. All data analyses were done in R v4.2.2 and graphs were created with the *ggplot2* package (R Core Team 2023).

## Results

### Detection likelihood

DNA of *G. campestris* was found in 20 of the 63 soil samples (31.75%). The species was discovered in six out of seven grasslands, including in two samples from the Åckelsta grassland, collected near the last known location where the species had last been observed eleven years ago, despite annual resurvey efforts. Our logistical models identified that both distance to the nearest individual of *G. campestris* and population characteristics affect *G. campestris*’ detection likelihood; in order of importance: 2021 population density (SES = 1.48) > nearest distance (|SES| = 1.27) > population size (SES = 0.78). Specifically, both declining distance to the nearest *G. campestris* individual (Model Chi^2^ = 14.00, p < 0.001) as well as increasing population size (Model Chi^2^ = 5.21, p = 0.02) had significant direct, positive effects on the species’ detection likelihood. Increasing population density in 2021 significantly elevated the species’ detection likelihood (Model Chi^2^ = 12.37, p < 0.001), though the latter was unaffected by population density in 2022 (Model Chi^2^ = 1.29, p = 0.26). When investigating the additive effects of population size on *G. campestris* detection likelihood, population size did significantly improve the distance model (Chi^2^ = 3.96, p = 0.04), translating in significant effect sizes for both distance (SES = −1.16, p = 0.002) and population size (SES = 0.73, p = 0.03). The additive effect of population size did improve the 2021 population density model (Chi^2^ = 10.41, p = 0.001), with a positive population density effect (SES = 1.20, p = 0.02), but no significant effect of population size (SES = 0.60, p = 0.06).

Depending on if RRA filtering was applied, *G. campestris* DNA was only detected in seven out of 63 soil samples (11.11%), and only found in three grasslands. Modeling results were, however, near-identical with direct positive significant impacts on detection likelihood by declining distance to the nearest *G. campestris* individual, increasing 2021 population density and increasing population size, and significant additive effects beyond distance of both population density as well as population size (for full model details, see Table S3 in Supplementary Information). Rather than distance to the nearest individual being most important, SES across different models suggested that population characteristics were more important in explaining *G. campestris’* detection likelihood. Finally, RRA filtering caused the likelihood of detecting *G. campestris* in soil cores taken closest to the last known location of the population, at the highest population densities and in large populations to decline with ca. 40%, 40% and 20%, respectively. Moreover, instead of more gradual changes in detection likelihood, RRA filtering led to rapid exponential declines in detection likelihood with declining population size and increasing distances.

### Community representation

The 63 eDNA soil samples and vegetation surveys together yielded a total of 141 unique plant species. A total of 67 species were identified using eDNA, of which three were exclusive to the eDNA analysis: *Agrostis vinealis, Stellaria alsine and Viola riviniana*. Hence, 46.43% of the observed plant species were uncovered using eDNA. RRA filtering removed 0.13% of all ASV reads, which equally represented the removal of 446 (out of 986; 45.23%) species × sample occurrences. Moreover, 8 observed species were lost from the original eDNA species × community matrix (17 unique species × sample occurrences; 1.72 %), including *A. vinealis*. eDNA species richness in a core sample decreased significantly due to RRA-filtering of the original eDNA data matrix (paired t-test: mean of 15.65 species vs. 8.57 species; t = 16.72, p < 0.001), yet RRA-filtered eDNA species richness was still correlated with eDNA species richness based on all presence/absences (r_Pearson_ = 0.39, p = 0.001). A full species list is available in Table S5 in Supplementary Materials, listing which species was uncovered by either method, or both.

Using the original full raw read sequence data matrix, eDNA samples contained significantly more species than the extant plant community at the 10 × 10 cm^2^ plot size (SES = −0.34; p < 0.001), whereas eDNA samples significantly contained less species than the extant community for the remaining plot sizes (SES_25cm_ = 0.19 < SES_50cm_ = 0.47 < SES_100cm_ = 0.71; p < 0.001). After RRA filtering, eDNA samples consistently and significantly contained less species than the extant community at each plot size (p < 0.001), yet with a steadily increasing difference in species richness with plot size (SES_10cm_ = 0.26 < SES_25cm_ = 0.79 < SES_50cm_ = 1.07 < SES_100cm_ = 1.31). In both instances, these patterns seem mainly driven by significantly increasing numbers of species unique to the extant communities (red boxplots in Fig. 4), i.e. species not detected in the eDNA sample. At the same time, species numbers detected in both the eDNA sample and the extant community (recorded by the traditional survey) significantly increased with plot size (blue boxplots in Fig. 4). This may be a result of species present at larger plot sizes were successfully uncovered by the eDNA analysis, as these species were actually being detected in the extant community. Consequently, a significant, gradual decrease in species numbers only detected by eDNA was also detected (green boxplots in Fig. 4). Still, RRA filtering caused less shared species to be uncovered, implying that this common data curation technique excluded many species confirmed to be present at the sites.

Compositional similarity between an eDNA core sample and its corresponding plot’s plant community was significantly affected by the choice of metric, with the Raup-Crick metric significantly reduced by ca. 2/3 compared to the typically-used Bray-Curtis metric, irrespective whether the eDNA data were RRA filtered or not (Fig. 3a). The suspected species richness bias underlying Bray-Curtis overestimation was underlined when RRA filtering was applied on the eDNA matrix. Species removal from the eDNA matrix significantly increased Bray-Curtis compositional similarity (t_Paired_ = −8.87,p < 0.001), as opposed to significantly reducing Raup-Crick similarity (t_Paired_ = 9.37, p < 0.001), which corrects for differences in species richness (Fig. 3a).

The Raup-Crick compositional similarity between an eDNA core (after RRA filtering) and a plot’s plant community was highest at the 10 × 10 cm^2^ plot size (mean 30 ± 23%; Fig. 3b), which was significantly higher than eDNAs compositional similarity with the plant community at all other plot sizes (SES_25cm|50cm_ = −0.10, SES_100cm_ = −0.05; p < 0.001). While eDNA compositional similarity was lowest and did not differ between the 25 cm and 50 cm plot sizes (21 ± 20% – 18 ± 19%, respectively). eDNA compositional similarity at 100 cm plot size proved to be significantly higher than similarity at 25 cm and 50 cm plot sizes (25 ± 23%, p < 0.01).

### Capturing and predicting plant community variation

eDNA species richness based on the full presence/absence data matrix was unable to predict species richness observed in the extant plant community at any plot size (Fig. 5 a-d). After applying a <0.05% RRA filter, eDNA species richness could significantly predict species richness of the extant plant community at 50 cm and 100 cm plot sizes (p = 0.01 and 0.003, respectively), but not at the small plot sizes (10 cm and 25 cm; p = 0.17 and 0.12, respectively; Fig. 5 e-h). eDNA samples could thus reliably discern between vegetation plots with low vs. high species richness in plot sizes commonly applied in grassland plant inventories when low read abundances were removed from the observed eDNA matrix.

All NMDS ordinations provided stable two-axes solutions with stress always ≤ 0.19, implying a good goodness-of-fit relative to the original sample positions in the multidimensional ordination space. For the full eDNA data matrix, the first vegetation NMDS axis was not related to the first eDNA NMDS axis at the 10 cm and 25 cm plot sizes (5.21 - 3.43% Deviance Explained; SES = 0.23 - 0.18, respectively; p > 0.05), while a significant relation was found for the 50 cm and 100 cm plot sizes (SES = 0.26 and 0.36, p = 0.04 and 0.004, respectively) even though with only 6.63 and 13.00% Deviance Explained, respectively. The second vegetation NMDS axis was significantly related to the second eDNA NMDS axis at all plot sizes, with %Deviance Explained increasing from 17.32% (SES = −0.42; p <0.001; 10 × 10 cm^2^) to 56.80% (SES = 0.75; p <0.001; 100 × 100 cm^2^). For the 0.05% RRA filtered eDNA data matrix, all general linear models at each plot size signaled that both eDNA NMDS axes could consistently and significantly predict their vegetation NMDS counterparts, yet with increasing accuracy as the % explained Deviance increased in absolute terms by 19% (NMDS1) and 43% (NMDS2) from the 10 × 10 cm^2^ to the 100 × 100 cm^2^ plot size. Plant community compositional variation was thus best explained by eDNA cores samples at the 100 × 100 cm^2^ plot size: the first eDNA NMDS axis significantly predicted compositional variation along the first vegetation NMDS axis (SES = −0.74; p < 0.001) with 54.84% of the deviance explained. The second eDNA NMDS axis also significantly predicted 69.68% of the compositional variation along the first vegetation NMDS axis (SES = −0.83; p < 0.001). This was also strikingly visible in both NMDS scatterplots (Fig. 6a, b), via comparable clustering patterns of the seven semi-natural grassland sites. For full results on NMDS ordinations, see Table S4 in Supplementary Materials.

## Discussion

Our study clearly indicates that soil-based eDNA metabarcoding may indeed deliver on its potential as a stand-alone tool for the discovery of rare plant species and plant community compositional change, aligning with the few studies available in natural terrestrial ecosystems today (Yoccoz et al. 2012, Edwards et al. 2018, Carrasco-Puga et al. 2021, Barnes et al. 2022, Ariza et al. 2023, 2024). Moreover, it is the first study which proves eDNA metabarcoding can detect a single rare species as well as comparatively subtle plant community change in highly diverse semi-natural grassland communities (up to 60 species per m^2^), which are the most diverse globally at small scales (Wilson et al. 2012), i.e. precisely at those small scales relevant to sampling design. Our findings also convincingly illustrate that sampling design, data curation and analytical parameters need to be specifically tailored to the planned objectives or research questions to ensure that correct ecological or practical inference can reliably be drawn based on the signals picked up in plant species and community data generated via soil-based eDNA metabarcoding.

### Detection likelihood of rare species

Our results confirm that soil eDNA metabarcoding can repeatedly detect the presence of a single rare plant species amongst 144 recorded grassland species (Carrasco-Puga et al. 2021, but see Hartvig et al. 2021). Even given significant site-specific variation in presence and abundance of the species, robust detection patterns emerged. Proximity to *G. campestris* and population characteristics both shaped the likelihood of detecting *G. campestris* DNA in a soil core (cf. Hartvig et al. 2021). Detection likelihood is indeed expected to decline as populations become smaller and more distant from the point of sampling, mirroring either distance-decaying DNA inputs (via e.g. pollen or propagules (Nakanishi et al. 2021, Nathan & Muller-Landau 2000) and leaf or root tissues (Levy-Booth et al. 2007, Yoccoz et al. 2012)) or other spatially clustered belowground phenomena such as soil seed banks (Plue & Hermy 2012), which *G. campestris* can build (Lennartsson & Oostermeijer 2001). Still, in stark contrast to Hartvig et al. (2021) who only found DNA at distances less than 10 cm from individual orchid plants, traces of *G. campestris* DNA are picked up over significant distances away from the populations, with a 25% detection likelihood at 25m (10% at 50 meters) from the closest individual, likely reflecting a combination of DNA distribution processes such as the long tails typical of seed and pollen dispersal events (Bullock & Clarke 2000, Austerlitz et al. 2004), leaf litter transport (Phillipson et al. 1975), or the persistence of local inter- and extracellular DNA of plants no longer present locally. Regardless of the source, with a more than reasonable effort of only nine soil cores per grassland, eDNA established the species presence in six out of seven grasslands.

However, to be of practical use for further conservation or restoration interventions, the eDNA signal has to be accurately interpreted. In that light, the observation that population density in 2022, the year after soil core sampling, had no significant impact on *G. campestris* detection likelihood seems crucial. This suggests that the eDNA signal is at least dominated by recently shed DNA, hence reflecting the presence of living *G. campestris* individuals, a notion also supported by the long DNA fragments amplified in this study (cf. Jo et al. 2017). Nevertheless, evidence suggests that the detected signal integrates both recently shed DNA and older DNA originating from seed banks, partially decomposed plant material, pollen and extracellular DNA retained in the soil (Carini et al. 2016, Ariza et al. 2023). Consequently, the eDNA signal should not be interpreted solely as a snapshot of the contemporary aboveground vegetation. On the one hand, *G. campestris* DNA was picked up in two out of nine soil cores in a grassland (*Åckelsta*) where the species’ presence had not been visually confirmed for over a decade in spite of targeted, annual ocular monitoring efforts. This observation thus clearly suggests that at least part of the detected signal may originate from long-term DNA reservoirs in the soil (e.g. extracellular, Carini et al. 2016; seed bank, Ariza et al. 2023), although the exact age and source of the detected DNA cannot be determined from the present study. On the other hand, in two grasslands (*Östra Syninge* and *Snedbacken*), population size over the past eight years averaged around one individual annually, suggesting highly limited recent DNA inputs of the small-statured, slender grassland biennial. Consequently, this dual nature of the eDNA signal – combining both contemporary and legacy DNA – implies caution when interpreting detections as certain evidence of extant populations. Still, even if legacy DNA were to dominate the signal, the eDNA signal identifies sites with recent historical occupancy, the presence of persistent soil seed banks or otherwise overlooked populations that warrant targeted field verification. More generally, the benefits for conservation and restoration of applying soil eDNA metabarcoding may thus lie not only in detecting currently visible populations, but also in revealing ecological legacies that are difficult to capture through conventional surveys, yet which may justify and rationalize targeted conservation and restoration efforts.

The discovery of DNA of the species across spatial and temporal gradients using eDNA metabarcoding is an impressive feat, given the species’ small-stature, biennial lifespan, low biomass production and low seed production (Lennartsson & Oostermeijer 2001). This realisation is even more impressive given the large amounts of fresh DNA shed by the numerous other grassland species, which co-exist at small spatial scales (Wilson et al. 2012), in highly biologically active semi-natural grassland soils (Siebert et al. 2019), next to methodological repeated subsampling (core sample > 5g soil subsample for DNA extraction) and further technological bottlenecks (e.g. choice of primer or bioinformatic pipeline; sequencing depth) (Deagle et al. 2019, Barnes et al. 2022, Luo et al. 2023), potentially diluting the signal of *G. campestris*. Still, a point of concern is the strong observed reduction in detection likelihood following frequently applied RRA filtering, highlighting the trade-off between (conservative) contamination control and information loss. As noted, *G. campestris* likely only produces limited DNA amounts due to small population sizes, limited above-and belowground biomass production given its small stature and biennial lifespan (Ulrich et al. 2010, Edwards et al. 2018). Thus, filtering out low-abundance reads, while often motivated by concerns about false positives, may disproportionately remove true positives for rare species such as *G. campestris*. Our results therefore clearly caution against applying uniform abundance filtering thresholds in studies or applications using eDNA metabarcoding to target rare species. Instead, filtering strategies should be tailored to objectives: conservative filtering risks undermining the very goals of early detection, red-list monitoring, or restoration planning (Hartvig et al. 2021, Sepp et al. 2021).

### Community composition and compositional change

eDNA metabarcoding captured a substantial subset of the extant semi-natural grassland communities. Across all 63 plots, eDNA metabarcoding succeeded to identify ca. 50% of traditionally surveyed plant species, a level of taxonomic overlap closely aligned with earlier reports for soil-based eDNA studies in other less species rich natural terrestrial ecosystems (e.g. ~60%, tundra, Edwards et al. 2018; ~50%, desert, Carrasco-Puga et al. 2021; ~60%, coniferous forest, Ariza et al. 2023). This level of consistent compositional overlap across habitat types suggests that the only partial recovery of present species seems a general characteristic of soil eDNA metabarcoding at its current technological state-of-the-art and bottlenecks (Luo et al. 2023, Barnes et al. 2022), rather than a limitation imposed by the high plant diversity in semi-natural grasslands (Wilson et al. 2012).

Still, given (1) this baseline level of taxonomic overlap, (2) the observed distance-decay detection patterns in *G. campestris* and (3) the ca. 25% probability of failing to detect a species even when sampling in its immediate vicinity (Fig. 2), it is unsurprising that a single soil core did not provide a full account of a plot’s plant community across plot sizes, neither in terms of its richness (Fig. 4, 5) nor its composition (Fig. 3b). Moreover, the low, species-richness corrected Raup–Crick similarities suggest that the observed mismatch is not driven by richness differences alone, but by differential species discovery by either method (cf. Carrasco-Puga et al. 2021). Partitioning species richness into its shared, vegetation-only, and eDNA-only cohorts provides valuable insight into this pattern. Across all plot sizes, a significant share of species was detected exclusively by eDNA, highlighting that soil eDNA samples capture species not visible in aboveground surveys at any given plot size. Such detections may include species missed during vegetation inventories, but are more likely to reflect dormant life-cycle stages such as seeds and belowground organs, or persistent intercellular (e.g. pollen, roots) and extracellular DNA from species formerly present or located just outside the sampled plot (cf. Hiiesalu et al. 2012). The number of shared species and eDNA-only species indeed significantly increased and decreased with increasing plot size, respectively, underscoring that numerous species uncovered only by eDNA are species actually present within the local species pool surrounding a sampled plot, reaffirming the highly local nature of an eDNA signal (cf. Hartvig et al. 2021, Ariza et al. 2023). Finally, RRA-filtering magnified species partitioning patterns among cohorts, again asserting this procedure may be overly conservative in removing truthful, and thus ecologically valuable information, as increasingly less species were considered shared or uncovered by eDNA, notwithstanding these species actual presence. In sum, eDNA analysis of a single soil core does currently not yet function as a direct spatial analogue of aboveground vegetation across plot sizes, but rather as an integrative signal reflecting local species pools across fine spatial grains (cf. Hartvig et al. 2021). Therefore, at plot scales commonly used in semi-natural grassland surveys, composite sampling will be necessary for applications requiring accurate representation of plant community richness and composition, as applied for similar invisible, and highly spatially clustered belowground phenomena (Baker et al. 2009, Muukkonen et al. 2009, Plue & Hermy 2012).

**Fig. 2.**
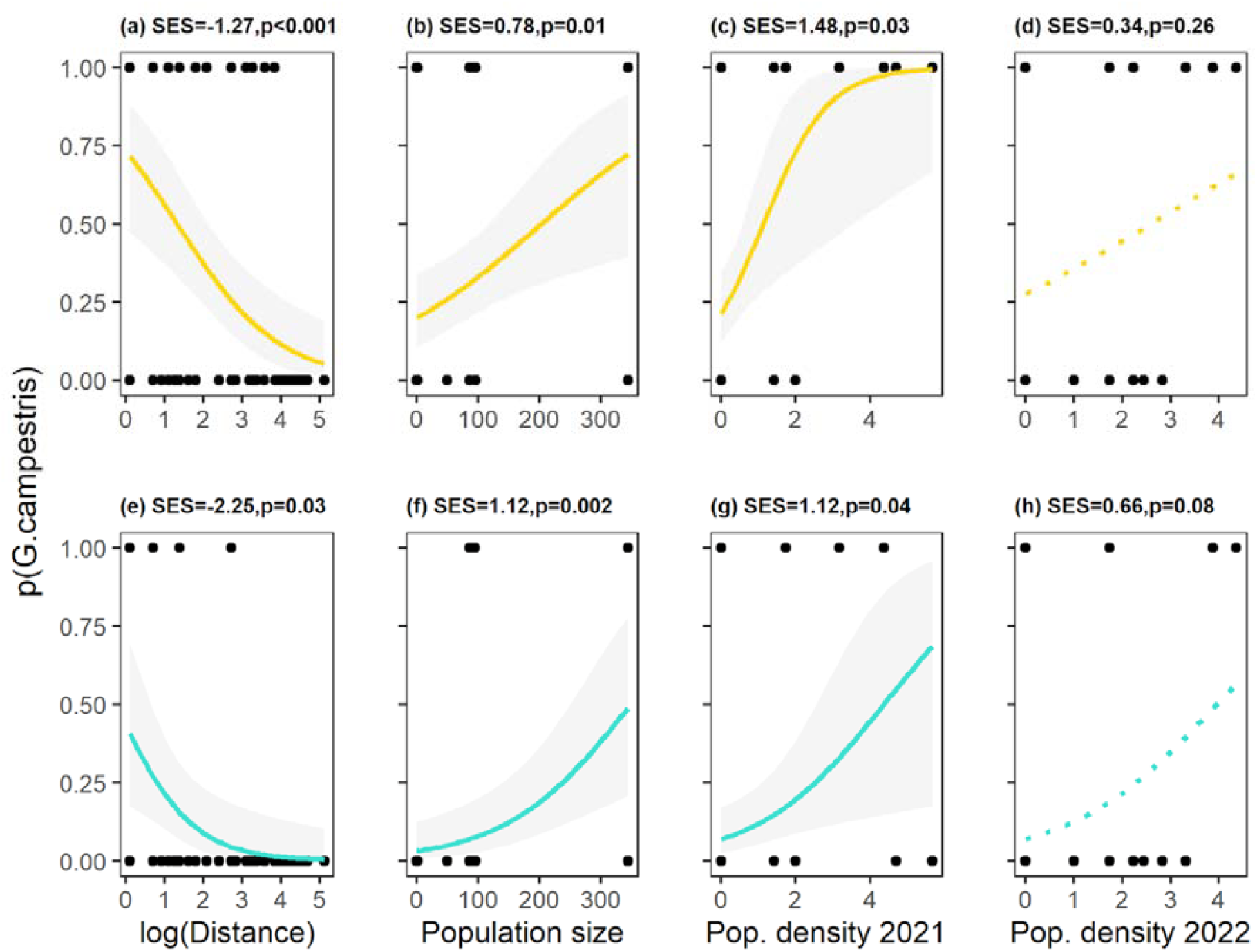
Direct effects of distance to the nearest individual (a, e), population size (b, f) and population density in 2021 (c, g) and 2022 (d, h) on the detection likelihood of the grassland plant specialist *Gentianella campestris* (*p(G. campestris)*) across 63 soil core samples, both for the full presence/absence data (top row, gold) and the RRA filtered presence/absence data (bottom row, turquoise). Standardized effect sizes (*SES*) and associated p-values are calculated via mixed effects logistic regressions using scaled predictor variables to enable comparison of the strength of the various predictor effects across models. Subplots (a) - (d) are ordered from right to left in decreasing importance of the predictors effect size on *G. campestris* detection likelihood.

**Fig. 3.**
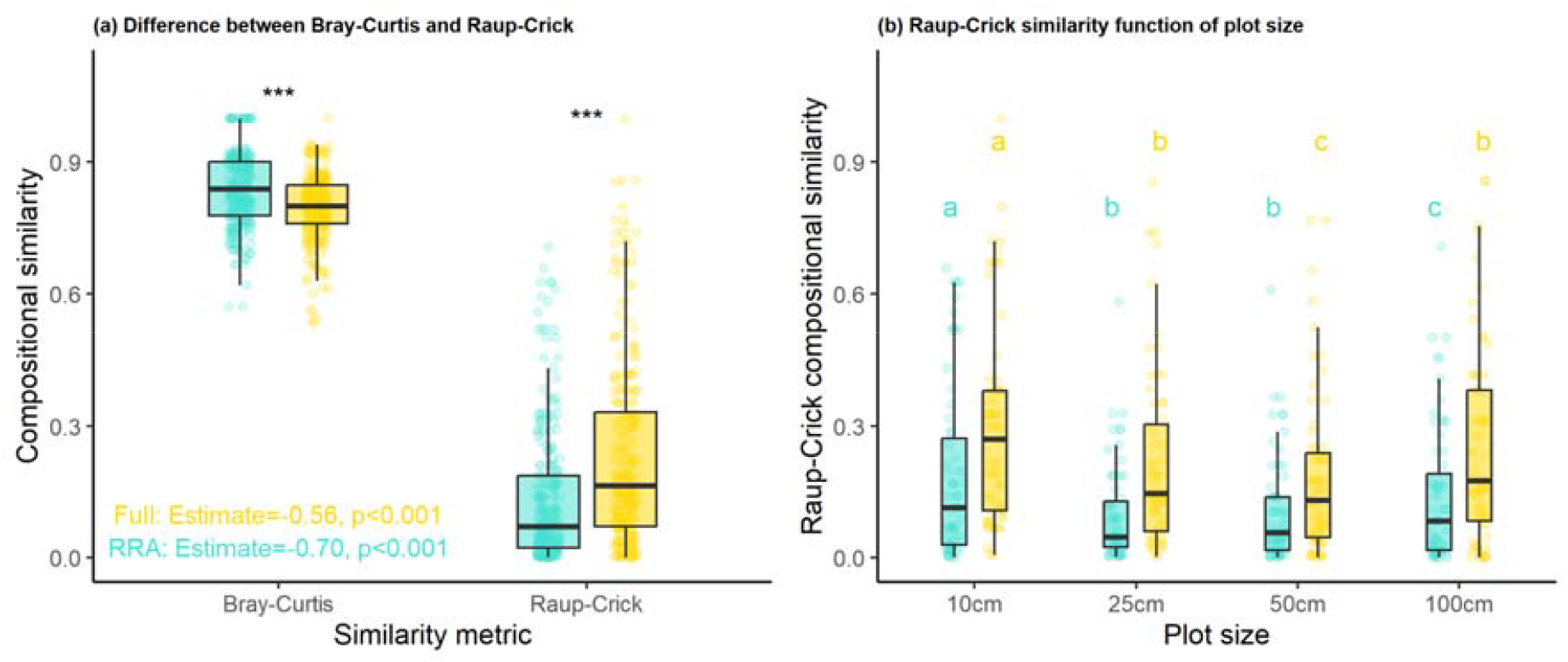
The impact of choice of similarity metric (a), plant inventory plot size (b) and data curation ((a) and (b)) on the compositional similarity between plant communities assessed via eDNA and traditional plot inventories. Colour coding visualizes the effect of data curation, i.e. when compositional similarity was calculated using either the full (gold) or 0.05% RRA-filtered eDNA dataset (turquoise).(a) Bray-Curtis similarity is significantly larger than Raup-Crick similarity, both when using the full (gold) and 0.05% RRA-filtered eDNA dataset (turquoise) to compare compositional similarity with the extant plant community. RRA filtering lead to significantly higher and lower similarity in the Bray-Curtis and Raup-Crick metric, respectively (***; p<0.001; paired t-test). (b) Homogeneous subsets

**Fig. 4.**
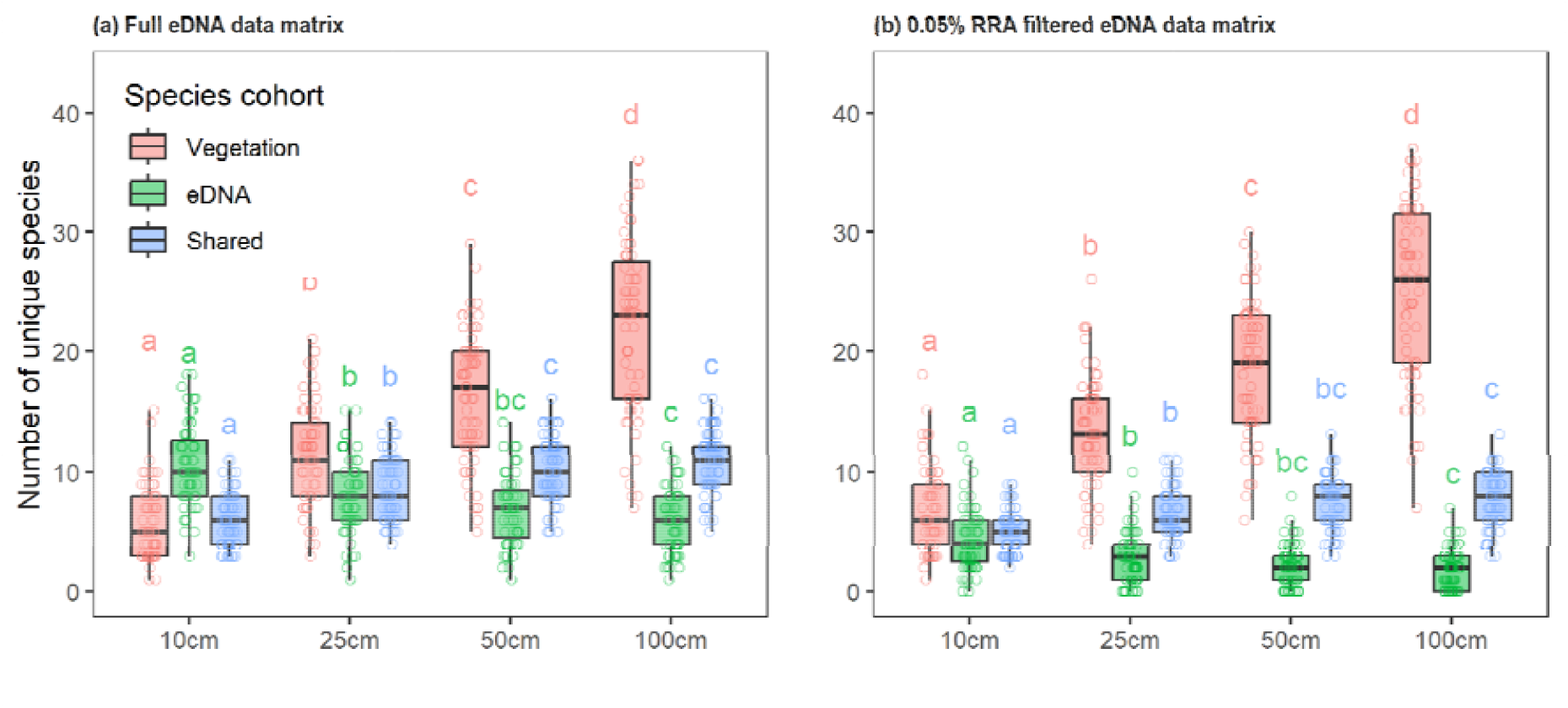
Partitioning of plant species uncovered by both eDNA and plant inventories at each sampling point as a function of plot size, before (a) and after (b) data curation by applying a 0.05% RRA filter on the full read abundance eDNA data. Species are partitioned as being uncovered: by the plant inventory only (red), by the eDNA assay only (green) or by both plant inventory and eDNA assay (blue). Homogeneous subsets identify significant shifts in species partitioning between inventory type with plot size.

**Fig. 5.**
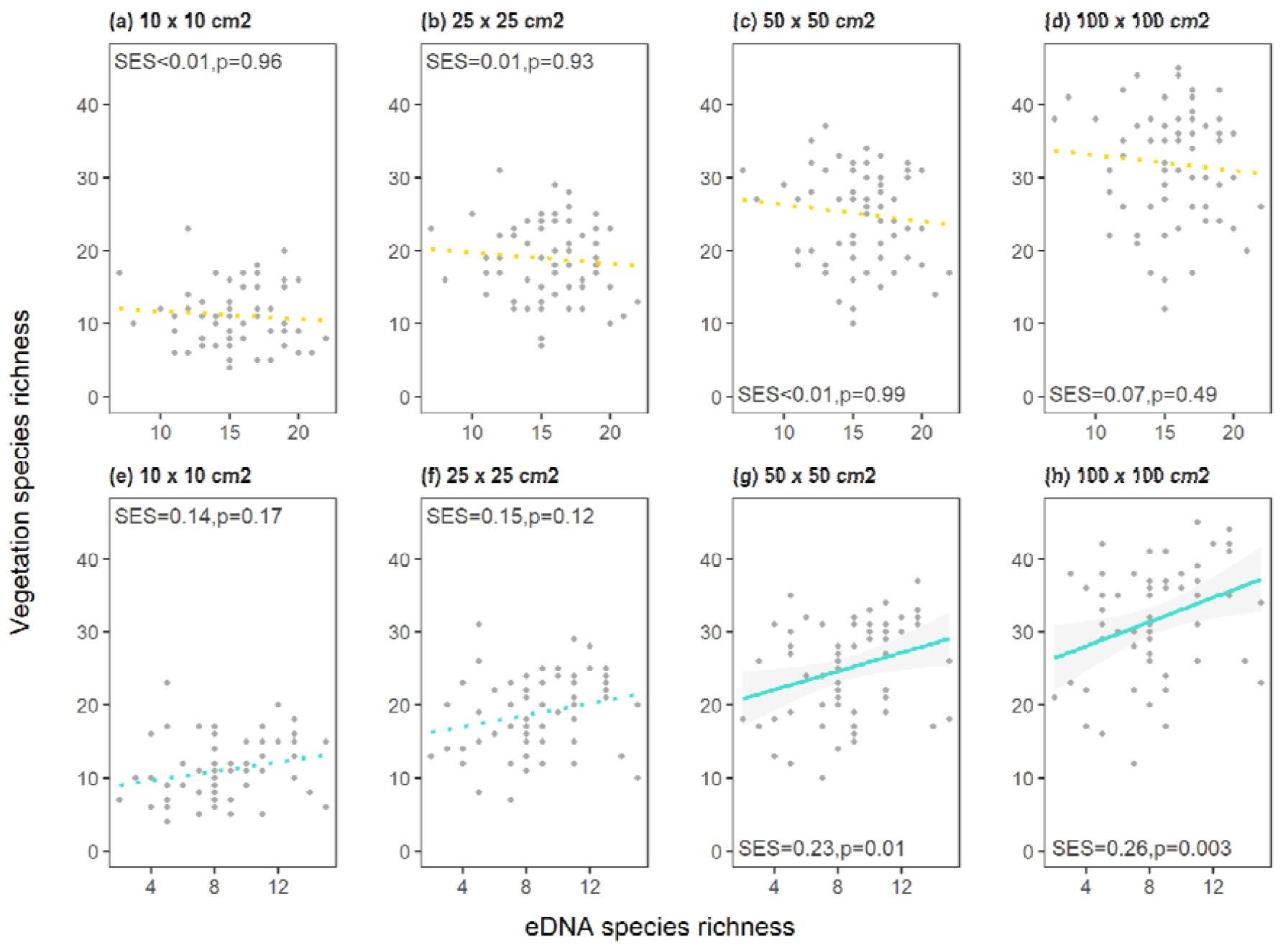
Relationships between species richness of the plant community identified using eDNA metabarcoding of 63 individual soil cores and quantified using plot-based inventories of different size (10 (a, e), 25 (b, f), 50 (c, g) and 100 cm (d, h)) nested within one another around each soil core. Relationships are presented for both the full presence/absence data (top row, gold) and the 0.05% RRA filtered presence/absence data (bottom row, turquoise), with full regression lines marking significant relationships at p < 0.05. Standardized effect sizes (*SES*) and associated p-values are calculated via general linear mixed models (including *site* as a random intercept) using scaled dependent variables to enable comparison of effect sizes across models.

**Fig. 6.**
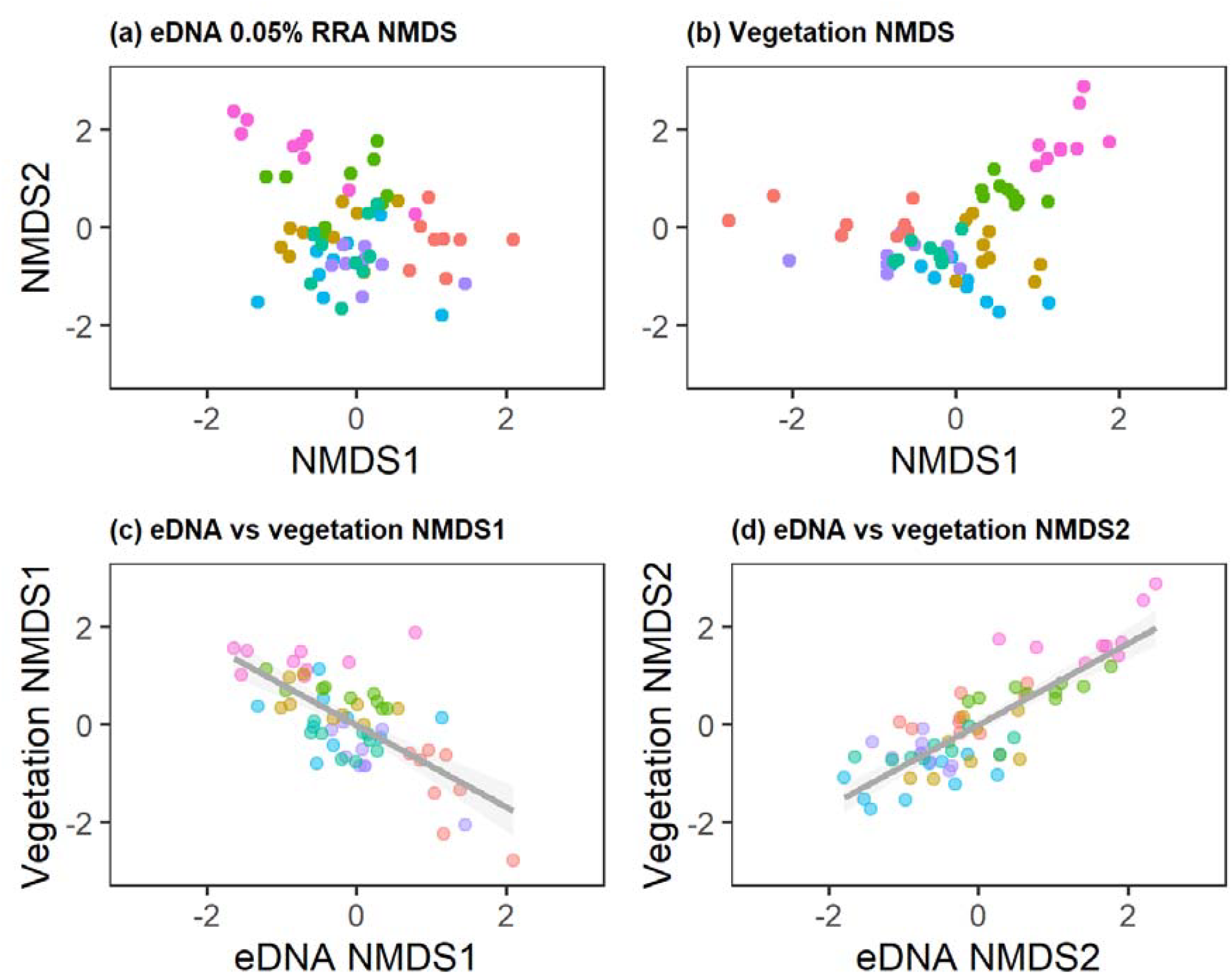
NMDS ordinations (based on Raup-Crick similarities) run separately on the 0.05% RRA filtered eDNA species × sample matrix (a) and observed extant plant communities within 100 × 100 cm^2^ plots centered around each of the 63 eDNA soil samples (b). Color coding identifies each of the seven semi-natural grasslands included in the study. Regression lines in subplots (c) and (d) highlight significant relations between community compositional gradients (NMDS axes) discovered separately in the eDNA and observed plant community data matrices.

Still, most plant ecological research rarely focuses on knowing the exact number and identity of every species at a given site (Vellend et al. 2008). Instead, we aim to detect changes in species richness and community composition in space and time, helping to pinpoint the environmental drivers which are shaping spatio-temporal plant community responses for conservation purposes. When removing local eDNA signal noise, richness estimates reliably tracked the variation in species richness at the 50 cm × 50 cm and 100 cm × 100 cm plot scales (cf. 40 m × 40 m plots in Barnes et al. 2022). Similarly, NMDS showed that eDNA-derived community composition not only mirrored subtle vegetation-based compositional gradients but quite accurately discriminated individual semi-natural grasslands (cf. Hartvig et al. 2021, Ariza et al. 2024). Combined, this implies that - using the sampling method described in this study - the species subset captured via eDNA metabarcoding both retains sufficient plant community structure to distinguish plots of low versus high diversity as well as identify the core ecological gradients shaping the spatial and environmental structure of semi-natural grassland communities, both within and across sampled semi-grasslands. This aligns with Vellend et al. (2008)’s argument demonstrating that subsets of species, even if incomplete (missing in some cases up to 80% of occurring species), can still reliably reflect patterns of variation in richness and composition if they include sufficient information from the broader community, i.e. if subsets are non-random and representative. As the Raup–Crick similarities confirm eDNA species assemblage to be a non-random, compositional subset of the observed community, the strong synchrony in conveying similar ecological information about spatial patterns of plant community turnover between eDNA metabarcoding and classical inventories therefore strongly suggests that eDNA metabarcoding indeed uncovers representative –and thus ecologically meaningful – subsets of species.

## Conclusion

The purpose of ecological surveys is to learn as much as we can with limited resources when, for example, mapping rare species distributions or quantifying changes in species richness or community composition by means of efficient, representative sampling strategies (cf. Vellend et al. 2008; Plue & Hermy 2012). eDNA metabarcoding is the latest addition to this sampling toolbox, with results favourably arguing for soil-based eDNA metabarcoding possibly becoming a successful stand alone tool at delivering a cost- and resource-effective means for surveying diverse terrestrial plant communities, pending that study objectives, sampling strategies and analytical choices are well-aligned. Our results highlight both the main practical consequences for sampling strategies and analytical pathways which arise when choosing between a focus on detailed, complete plant community inventories including rare species detection versus unveiling spatial and temporal patterns in plant communities. Therefore, we believe that soil-based eDNA metabarcoding may presently not always replace traditional surveys for exhaustive plot-level species inventories, yet it is already a potent tool in providing a scalable, observer-independent tool for detecting biodiversity patterns, monitoring compositional change, and thus effectively informing conservation and restoration efforts.

## Acknowledgements

Mikaela Boltenstern is acknowledged for her field- and lab-work assistance during the project. The authors would also like to acknowledge support of the National Genomics Infrastructure (NGI) / Uppsala Genome Center and UPPMAX for providing assistance in massive parallel sequencing and computational infrastructure. Work performed at NGI / Uppsala Genome Center has been funded by RFI / VR and Science for Life Laboratory, Sweden. The molecular biology analysis of the soil samples were performed by the laboratory at IVL Swedish Environmental Research Institute, Stockholm. The authors have no conflict-of-interest to declare.

## Data availability

DNA-sequence data from this project has been deposited in GenBank under BioProject No. PRJNA1465734 [https://www.ncbi.nlm.nih.gov/bioproject/PRJNA1465734]. Upon acceptance, data will be archived in an online Dryad digital repository (Plue & Töpel, 2026).

## Supplementary Information

**Table S1.**
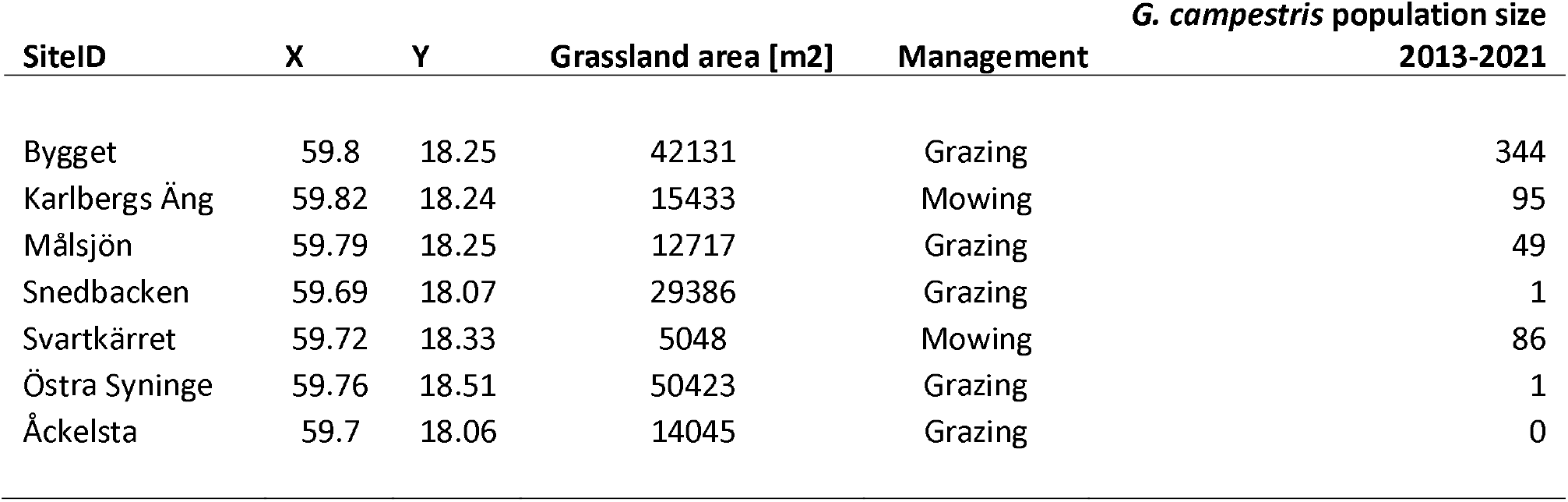
Basic characteristics of the seven semi-natural grasslands in Stockholm county (Sweden) in which 9 soil cores were collected for eDNA metabarcoding to identify.

**Table S2.**
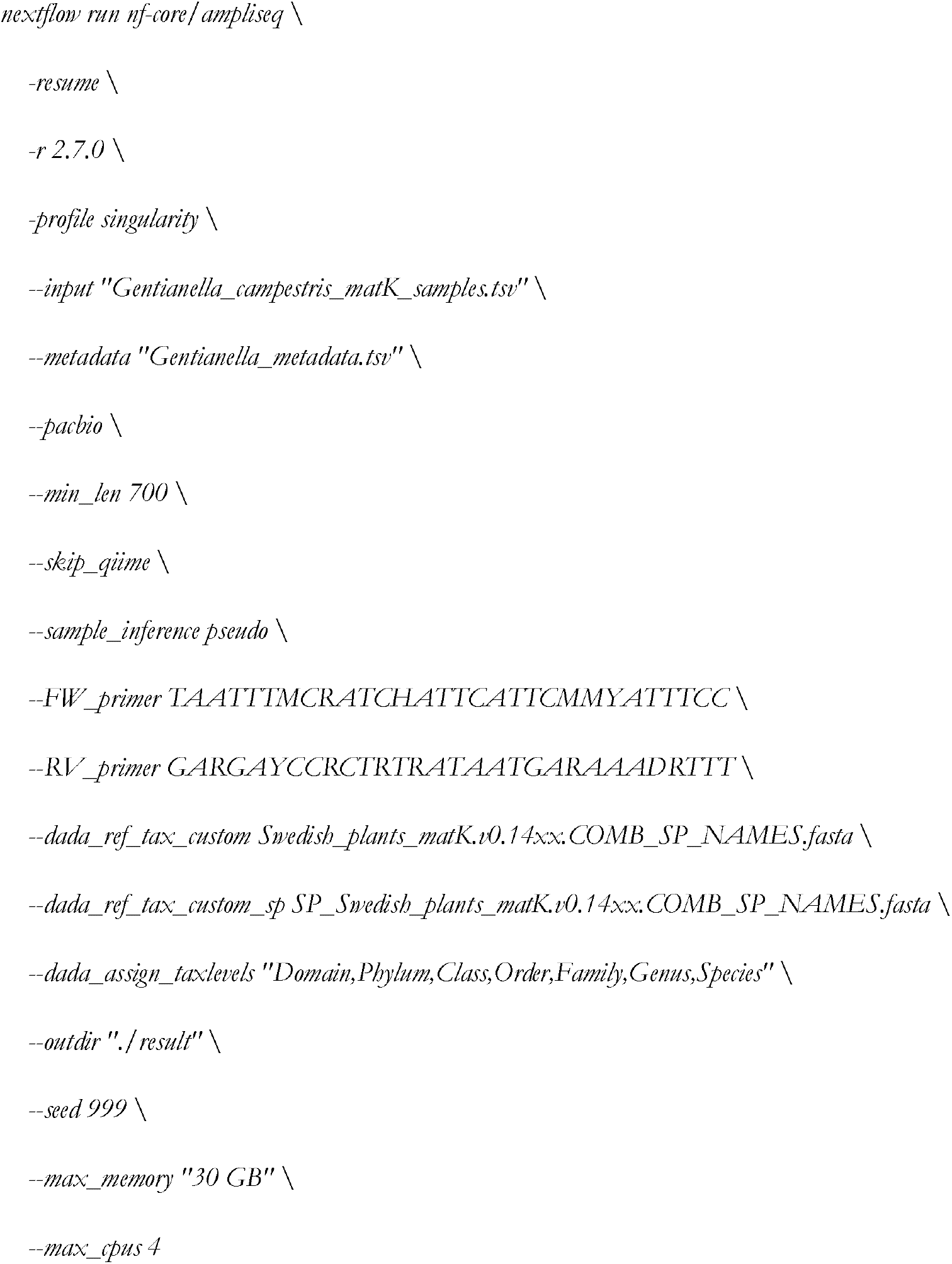
Parameters used in the bioinformatics pipeline used for taxonomic assignment of ASVs using the dada2 R package (Callahan et al. 2016).

**Table S3.**
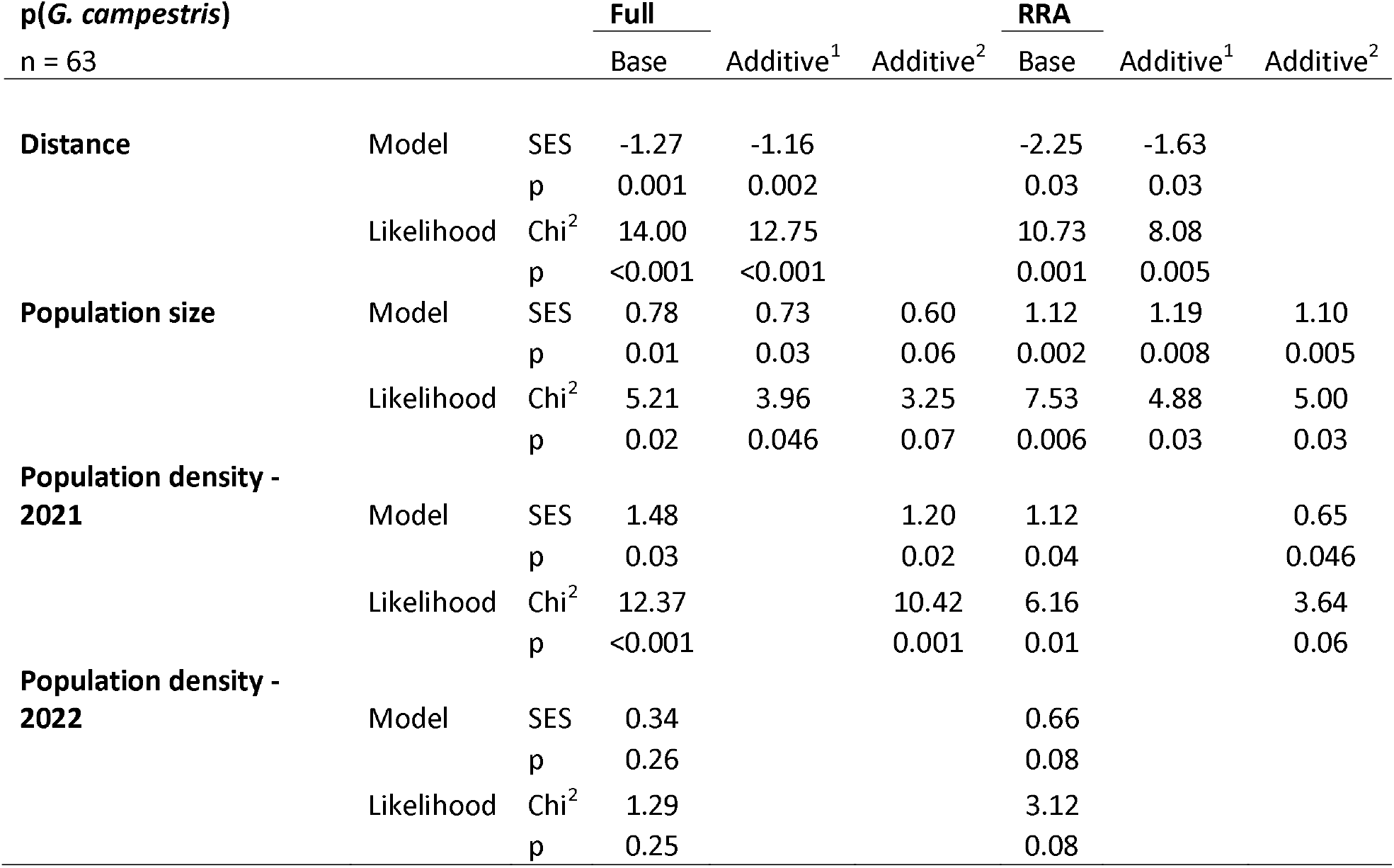
Detection likelihood of *Gentianella campestris* in an eDNA soil core sample in function of *G. campestris* population and sampling characteristics, modelled using logistic regression models, all containing *site* as a random model term. Base models contained a single scaled explanatory variable (returning Standardized Effect Sizes (SES) for comparability between models), using likelihood ratio tests to assess model improvement to a null model. In the additive models, we investigated the importance of *Distance* (1) and *2021 Population density* (2) by adding these variables separately to the base model containing *Population size* as explanatory variable, using likelihood ratio test to assess model improvement. This analysis was done both on the presence of *G. campestris* in an eDNA soil core sample before (full) and after (RRA) data curation, i.e. applying a <0.05% RRA omission filter on the raw ASV read abundance matrix. Distance was log-transformed whereas Population densities were square-root transformed for normality, prior to scaling.

**Table S4.**
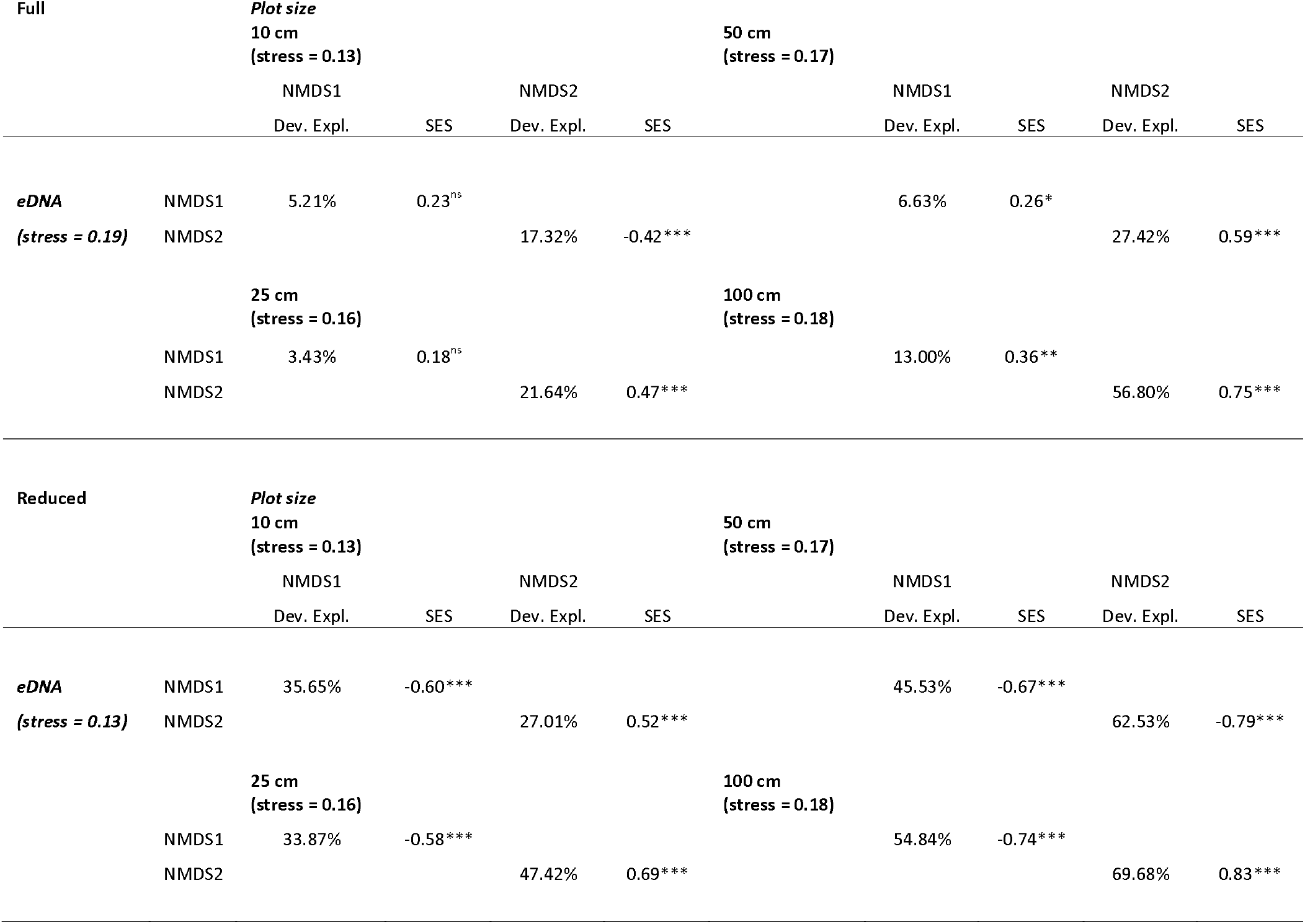
Explanatory power of eDNA plant community composition over extant plant community composition observed at different plot sizes (10, 25, 50 and 100 cm) both before (Full) and after (Reduced) data curation by removing all species observations (i.e. read counts) with less than 0.05% RRAs in the full eDNA species × sample matrix. Gradients in plant community composition were quantified using NMDS ordinations based on Raup-Crick similarities for eDNA and plant community data separately (final stress < 0.20 indicates good fit of the multidimensional data). The presence of significant relations between community compositional gradients discovered by eDNA and plot-based observations (NMDS axes) and their explanatory power (% Deviance Explained) were tested via general linear models between scaled NMDS axes. Scaling returned Standardized Effect Sizes (SES) enabling comparison between models.

**Table S5.**
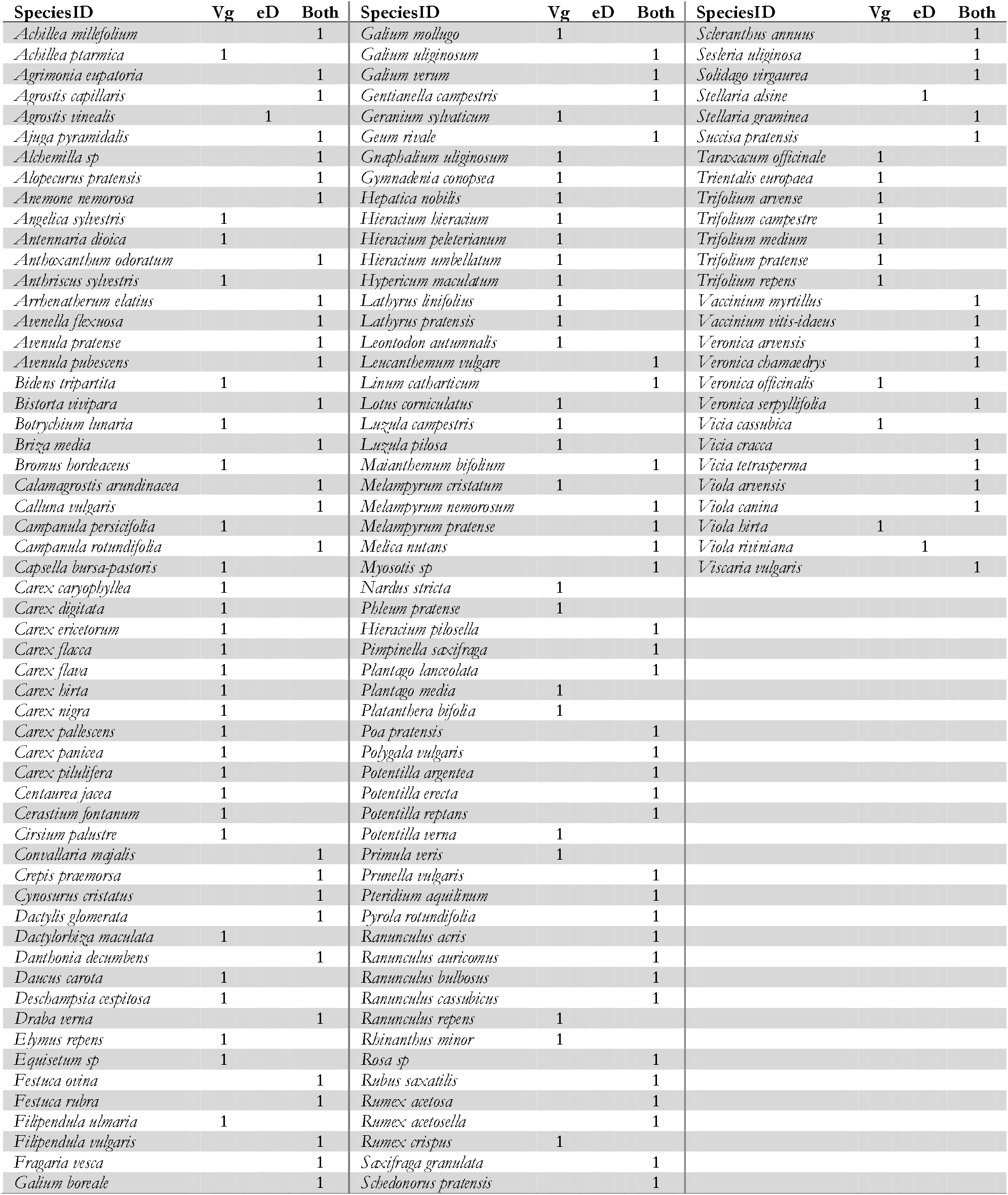
Complete list of the 141 plant species uncovered during this study across 7 semi-natural grasslands in Sweden, identifying whether each species was uncovered by either soil-based eDNA metabarcoding (eD), the vegetation surveys (Vg), or by both methods.

**Dataset S1 Soil core and *Gentianella campestris* population characteristics**

For each of 63 eDNA soil cores collected, this dataset lists: (1) the semi-natural grassland and (2) its size [m^2^] in which soil cores were collected, (3) the latitude and longitude per individual core, (4) the grassland management, (5) the distance of the soil core to the nearest and last known observation of *Gentianella campestris*, (6) 2021 and (7) 2022 *G. campestris* population density, i.e. the number of *G. campestris* individuals counted in a 2m radius surrounding each soil core, (8) mean population size of *G. campestris* in the last 8 years, (9) the last year *G. campestris was obs*erved in the semi-natural grassland and finally, if *G. campestris was det*ected by eDNA [0/1] before (10) and after (11) a 0.05% RRA filter was applied to the raw ASV read eDNA plant community matrix (Dataset S3).

**Dataset S2 Abundance-based plant community matrix**

Full 138 species × 252 plot-based plant community matrix. Four nested plant relevés of different plot sizes (10, 25, 50 and 100 cm) were performed centred around each of 63 individual soil cores collected for eDNA analysis, at a rate of 9 soil cores in each of 7 semi-natural grasslands in Stockholm county, Sweden.

**Dataset S3 ASV-based eDNA plant community matrix**

Full 67 species × 63 eDNA-based plant community matrix, after manual curation of the raw ASV abundance matrix as described in the main manuscript. Values represent the number of ASV reads assigned to a species via taxonomic bioinformatic pipelines (against libraries consisting of the vascular plant species present in Sweden), further manual data curation via BLASTn and removal of any ASVs based on suspected lab contamination.

